# Occurrence of cross-resistance and beta-lactam seesaw effect in glycopeptide, lipopeptide, and lipoglycopeptide-resistant MRSA correlates with membrane phosphatidylglycerol levels

**DOI:** 10.1101/671438

**Authors:** Kelly M. Hines, Tianwei Shen, Nate K. Ashford, Adam Waalkes, Kelsi Penewit, Elizabeth A. Holmes, Kathryn McLean, Stephen J. Salipante, Brian J. Werth, Libin Xu

## Abstract

Treatment of methicillin-resistant *Staphylococcus aureus* (MRSA) infections is challenging and is associated with high rates of therapeutic failure. The glycopeptide (GP) vancomycin and the lipopeptide (LP) daptomycin are still relied upon to manage invasive MRSA infections; however, resistance to these antibiotics has emerged and there is evidence of cross-resistance between them. It has been observed that the susceptibility of MRSA to beta-lactams increases as susceptibility to GPs and LPs decreases, a phenomenon termed the seesaw effect. Recent efforts to understand the mechanism underlying the seesaw effect have focused on the penicillin binding proteins (PBPs). However, while daptomycin resistance is largely mediated by remodeling of membrane lipid composition, the role of membrane lipids in producing cross-resistance and the seesaw effect has not yet been investigated. Here, we evaluate the lipid profiles, cross susceptibilities, and beta-lactam susceptibilities of a collection of isogenic MRSA strains selected against daptomycin, vancomycin or dalbavancin (a lipoglycopeptide; LGP) to assess the relationship between membrane composition, cross-resistance, and the seesaw effect. We found that modification of membrane composition occurs not only in daptomycin-selected strains, but also vancomycin- and dalbavancin-selected strains. Significantly, we observed that typically the levels of short-chain phosphatidylglycerols (PGs) negatively correlate with MICs of GP/LP/LGP and positively correlate with MIC of certain beta-lactams, the latter being dependent on the primary PBP target of the particular beta-lactam. Furthermore, changes to certain PGs with long-chain fatty acids correlate well with presence of the seesaw effect. These studies demonstrate a major association between membrane remodeling and the seesaw effect.

## INTRODUCTION

*Staphylococcus aureus* remains a serious public health concern that is responsible for nearly 120,000 infections and 20,000 deaths in the USA annually (1). Methicillin resistant *S. aureus* (MRSA) is particularly challenging to treat and nearly 40% of infected patients will fail therapy with the first-line antibiotic, vancomycin, a glycopeptide (GP) (2–7). Many factors contribute to poor outcomes in these patients including the emergence of reduced susceptibility phenotypes such as vancomycin intermediate *S. aureus* (VISA) and heterogeneous VISA (hVISA) (8, 9). Several alternatives for vancomycin are available, including the lipoglycopeptides (LGPs) dalbavancin, telavancin and oritavancin, and the lipopeptide (LP) daptomycin. However, the minimum inhibitory concentration (MIC) of vancomycin is positively correlated with MICs for LGPs and LPs, posing the risk of cross-resistance establishing among these therapies without the patient having been exposed to them (10, 11).

Many studies have observed that the susceptibility of MRSA to beta-lactams increases as the susceptibility to vancomycin or daptomycin decreases, a phenomenon now widely known as the beta-lactam seesaw effect (11–20). This phenomenon was first observed *in vitro* by Sieradzki *et al.* in laboratory-derived vancomycin-resistant MRSA strains (12–15). This dynamic between beta-lactams and peptide-based antibiotics has been shown to be therapeutically-relevant based on observations of the seesaw effect in clinical isolates (15, 17). While the exact mechanism leading to the seesaw effect is not known, some evidence suggests that it arises due to changes in the expression, abundance, or activity of penicillin binding proteins (PBPs) which coincides with glycopeptide and lipopeptide resistance (14, 21–26).

PBPs are the main machinery for cell wall synthesis and are the target of beta-lactam antibiotics. Methicillin resistance is mediated by the acquisition of the *mecA* gene encoding for an alternative PBP, PBP2a, that is not susceptible to traditional beta-lactam antibiotics targeting PBP1-PBP4. However, reduced susceptibility to vancomycin and daptomycin occurs through much different mechanisms. Vancomycin prevents crosslinking of the cell wall by binding to the peptidoglycan D-ala-D-ala residues and is most effective when the antibiotic can access the nascent peptidoglycan synthesized at the division septum (27, 28). VISA strains are often found to have more D-ala-D-ala binding sites and thick cell walls that prevent vancomycin from reaching the division septum (29–31). While several genes have been associated with the VISA/hVISA phenotype, thickening of the cell wall has been closely linked to mutations in the multi-component systems regulating cell wall metabolism, *yycFG* (or *walKR*) (32–34), and cell wall biosynthesis, *vraTSR* (35–38).

Daptomycin, a cationic lipopeptide, exerts bactericidal activity by targeting negatively-charged lipids in the membrane of *S. aureus*, followed by oligomerization and pore formation (39). One mechanism by which MRSA strains can effect daptomycin resistance is by decreasing the net-negative charge of the cell surface (40, 41). This occurs through the reduction of the major membrane lipid, phosphatidylglycerol (PG), which has a negatively-charged headgroup, or an increase in the modification of PGs with lysine (lysylPGs) to generate a positively-charged headgroup (40–44). Synthesis of lysylPGs is performed by mprF, and gain-of-function mutations or increased expression of the *mprF* gene is often observed in daptomycin resistance (45–49). Like VISA, daptomycin-resistant *S. aureus* may also have thickened cell walls but this is not a universal phenotype (50). Alternatively, increased D-alanylation of teichoic acid in the cell wall has been observed in daptomycin-resistant *S. aureus*, which could also contribute to the increased positive charge in the cell envelope (51, 52).

Dalbavancin is a recently-approved long-acting lipoglycopeptide that works in a manner similar to vancomycin. The lipid-like tail of dalbavancin improves its effectiveness over vancomycin by interacting with the membrane and stabilizing its interaction will the cell wall (53). Dalbavancin resistance has only recently been observed, but the presence of thickened cell walls similar to VISA and cross-resistance with daptomycin and vancomycin suggest that there may be overlap in their resistance mechanisms (54, 55). Clinical isolates with dalbavancin resistance have been found to have mutations in *yvqF* (or *vraT*) (54) and *rpoB* (55), which are also frequently observed in VISA (38, 56).

In many strains of MRSA, exposure to beta-lactam antibiotics can reverse some of the phenotypic changes associated with VISA or daptomycin-resistance. Exposure to the anti-MRSA cephalosporin, ceftaroline, reduced cell wall thickness and increased daptomycin binding in VISA with daptomycin resistance (57). Various conventional beta-lactams have also been effective at modulating cell surface charge to prevent daptomycin resistance in MRSA (58, 59). These benefits have led to the widespread exploration of combination therapies that use a beta-lactam to improve the effectiveness of daptomycin (60–62) and vancomycin (10, 63–65) or limit the potential for daptomycin resistance to emerge (58). Yet, despite the strong correlation between membrane lipids and daptomycin resistance, the role of lipids in mediating cross-resistance and the beta-lactam seesaw effect has not yet been explored.

In this study, we have assessed the occurrence of cross-resistance among GP/LP/LGP and the beta-lactam seesaw effect in MRSA strains that have been selected by *in vitro* serial passage for resistance to daptomycin, vancomycin and dalbavancin. The susceptibility profiles of these strains were evaluated against beta-lactams with varying specificities for the major PBPs of *S. aureus*, to explore variability in the emergence of the beta-lactam seesaw effect in strains with resistance to daptomycin, vancomycin or dalbavancin. We also evaluated the global lipid profiles of all study strains and found strong correlation between the presence of cross-resistance and the seesaw effect and the abundances of individual PG species. This work provides evidence that alterations in lipid metabolism contribute to the mechanism behind the cross-resistance among GP/LP/LGP and beta-lactam seesaw effect, and that modification of membrane lipid composition is not limited to daptomycin resistance alone.

## RESULTS

### In vitro selection for GP/LP/LGP resistance

Resistance to daptomycin, vancomycin and dalbavancin was selected on the MRSA N315 and JE2 genetic backgrounds using the serial passage method. Table 1 shows the antibiotic susceptibility and genetic profiles for the six *in vitro* selected mutant strains. In N315, mutants were generated with 8-fold reduced susceptibility to daptomycin (N315-Dap1), 128-fold reduced susceptibility to dalbavancin (N315-Dal0.5), and 16-fold reduced susceptibility to vancomycin (N315-Van8), based on minimum inhibitory concentration (MIC) to the respective antimicrobials. In JE2, mutants were generated with 16-fold reduced susceptibility to daptomycin (JE2-Dap2), 512-fold reduced susceptibility to dalbavancin (JE2-Dal2), and 8-fold reduced susceptibility against vancomycin (JE2-Van4).

**Table 1.**
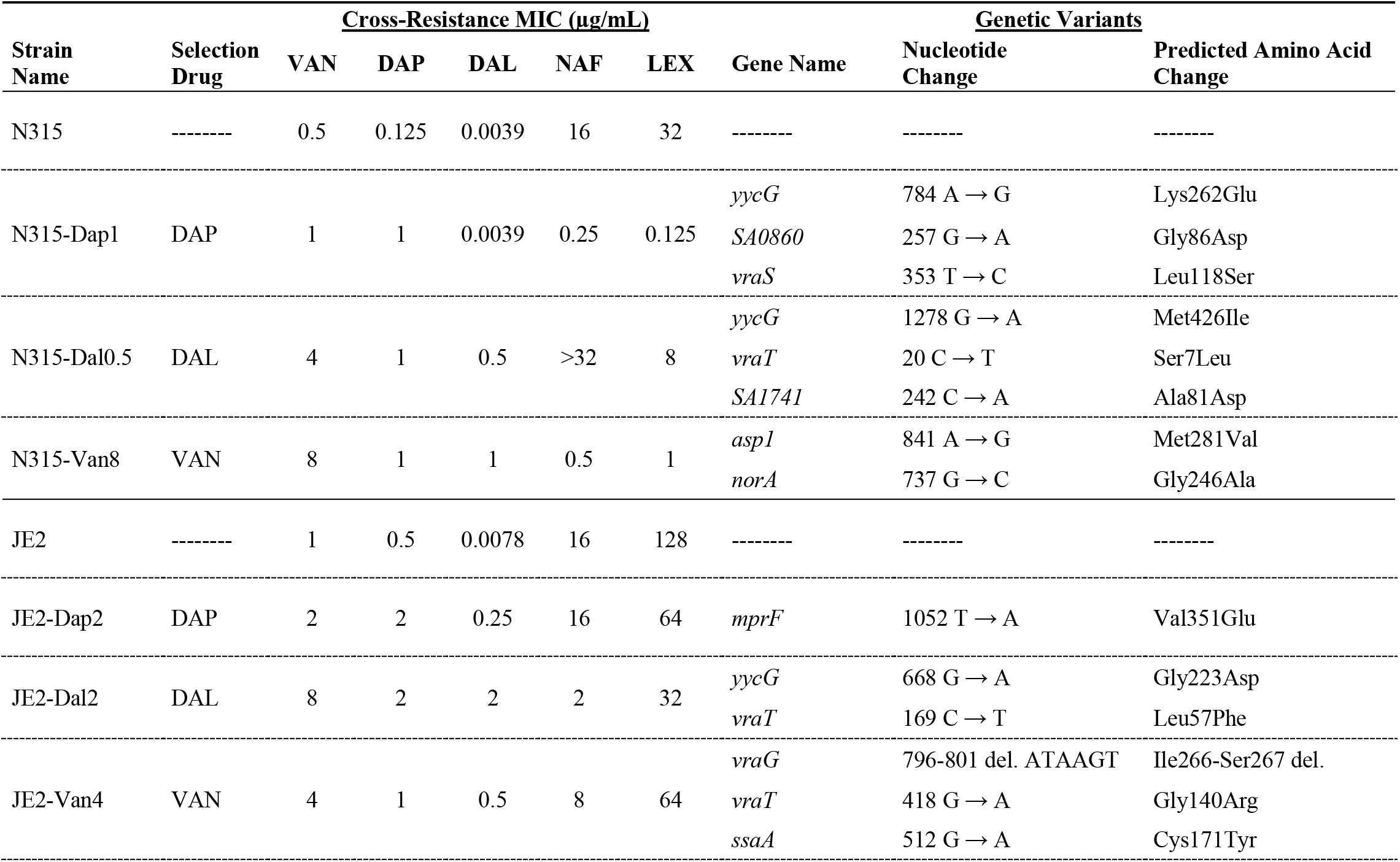
Strains used in this study and their antibiotic susceptibility and genetic profiles.

Whole genome sequencing (WGS) was performed on all selected strains to identify mutations which arose relative to the parental N315 or JE2 genotypes (Table 1). The most frequently mutated genes were *vraT* of the *vraTSR* operon (in N315-Dal0.5, JE2-Dal2, and JE2-Van4), and the histidine kinase *yycG* of the *yycGF* operon (in N315-Dap1, N315-Dal0.5, and JE2-Dal2). The histidine kinase *vraS* of the *vraTSR* system was also mutated in N315-Dap1. The genome of N315-Van8 additionally contained a variant in *asp1*, encoding accessory secretory system protein, and *norA*, a known contributor to quinolone resistance. A mutation in lysyl phosphatidylglycerol synthase *mprF* was identified exclusively in JE2-Dap2. JE2-Van4 contained a unique variant in a gene that encodes for secretory agent *ssaA*.

### GP/LP/LGP cross-resistance

Cross-resistance was assessed by MIC testing for these six strains, as shown in Table 1 and Figure 1. Serial passage with dalbavancin selected for resistance to vancomycin and daptomycin in both genetic backgrounds, with 8-fold reduced susceptibility against daptomycin in N315-Dal0.5, 4-fold reduced susceptibility against daptomycin in JE2-Dal2, and 8-fold reduced susceptibility against vancomycin in both N315-Dal0.5 and JE2-Dal2. In both genetic backgrounds, serial passage with vancomycin selected for cross-resistance to dalbavancin and decreased susceptibility to daptomycin. Serial passage with daptomycin selected for minor reductions to vancomycin susceptibility in JE2 and N315. However, JE2-Dap2 showed substantially increased resistance to dalbavancin with a 256-fold higher MIC relative to the parent JE2 strain, whereas no change in the dalbavancin MIC was observed in N315-Dap1. The lipoglycopeptide telavancin was also evaluated against the N315- and JE2-derived mutants (Supporting Information Table SI-1 and Figure 1). In N315 derivatives, the telavancin MIC were consistent with the trend of reduced susceptibility observed for dalbavancin. However, no meaningful change to telavancin MIC was observed in the JE2-derived strains.

**Figure 1.**
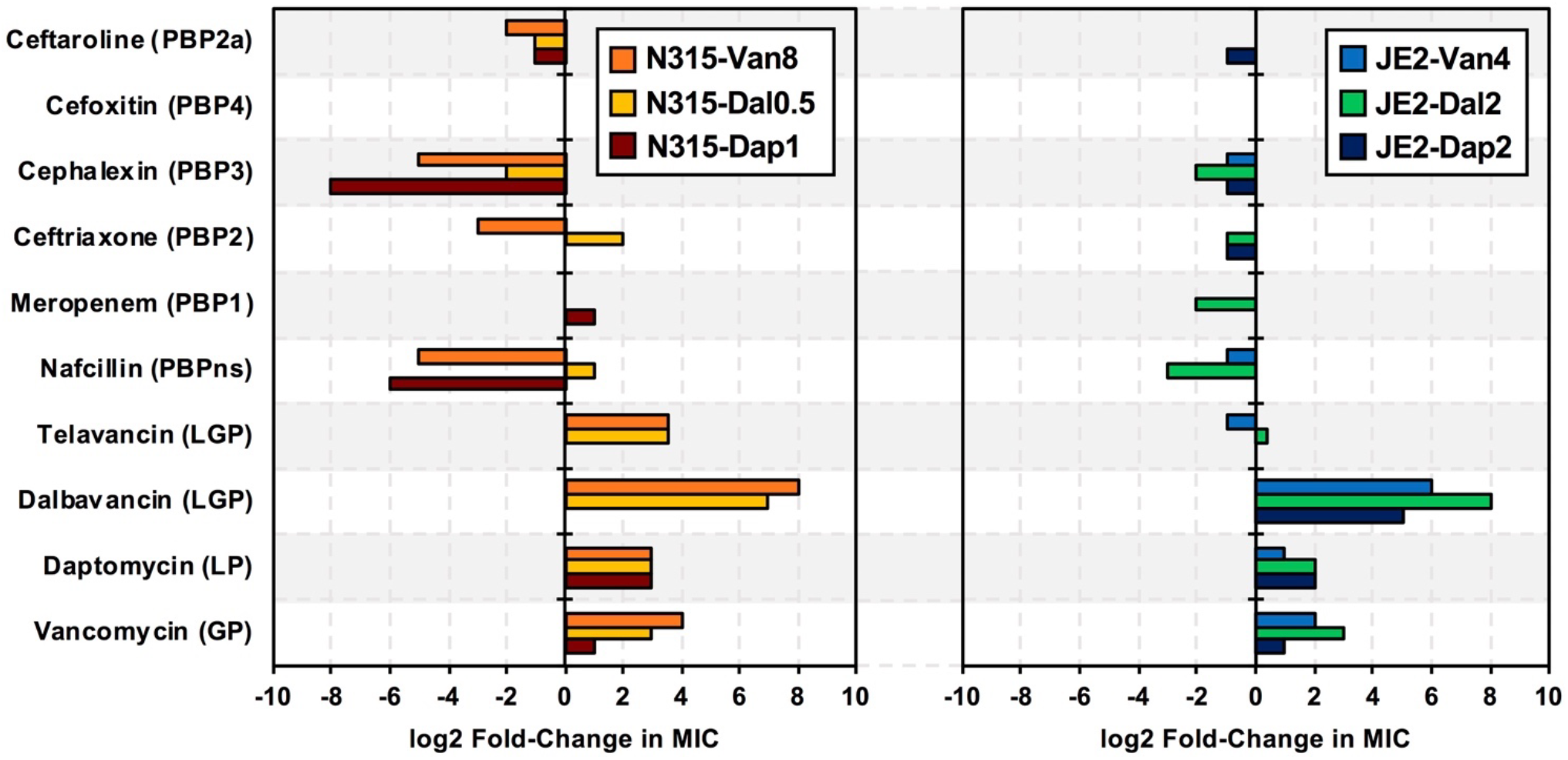
Susceptibility of A) N315-derived strains and B) JE2-derived strains against antibiotics targeting different penicillin binding proteins (PBPs) and peptide-based antibiotics (LGP, lipoglycopeptide; LP, lipopeptide; GP, glycopeptide). Log_2_ fold-change relative to the minimum inhibitory concentration (MIC) of the parent (N315 or JE2) strain.

### In vitro susceptibility to beta-lactams

The susceptibilities against a collection of beta-lactams with varied PBP specificities were determined for all study strains to assess the presence of the seesaw effect in a background of GP, LP, and LGP resistance. Table 1 shows the results from the susceptibility tests against the penicillin nafcillin, which has no PBP specificity. A strong seesaw effect with nafcillin was observed for N315-Dap1 and N315-Van8, for which nafcillin MIC decreased 64- and 32-fold, respectively. However, the MIC of nafcillin against N315-Dal0.5 was 2-fold higher than that of the N315-parent strain. In JE2, the largest fold-change difference in nafcillin MIC was observed for JE2-Dal2, which had an 8-fold reduction in MIC relative to the parent JE2 strain. The nafcillin MIC against JE2-Van4 decreased by 2-fold, whereas no change was observed against JE2-Dap2.

All study strains demonstrated seesaw effect with the PBP3-specific cephalosporin cephalexin (Table 1), but the extent of the susceptibility change varied substantially among strains. N315-Van8 and N315-Dap1 had substantially increased susceptibility to cephalexin, with 32- and 256-fold decreases in cephalexin MICs, respectively. The changes in the cephalexin MIC for the other study strains were much smaller, ranging from 2-4-fold lower than the parent strains.

Results for other beta-lactams are shown graphically in Figure 1 and in Supporting Information Table SI-1. All selected N315 strains evidenced 2-to 4-fold increased susceptibility to ceftaroline (PBP2a-specific), but only N315-Van8 displayed increased susceptibility to ceftriaxone (PBP2-specific). Conversely, N315-Dal0.5 showed a 4-fold increase in MIC against ceftriaxone. The MICs of meropenem (PBP1-specific) and cefoxitin (PBP4-specific) against the N315-derived mutants were unchanged relative to the parent strain, except for a 2-fold increase in the meropenem MIC for N315-Dap1.

Among JE2 strains, the JE2-Dal2 strain showed increased susceptibility to ceftriaxone (2-fold), cephalexin (4-fold), and meropenem (4-fold). JE2-Dap2 also showed 2-fold increased susceptibility to ceftriaxone, cephalexin, and ceftaroline. JE2-Van4 only displayed increased susceptibility towards nafcillin. The MIC of cefoxitin against all JE2-derived mutant strains was unchanged relative to the parent JE2 strain.

Collectively, these results reveal the presence of a seesaw effect for several beta-lactams in N315 and JE2-derived strains with resistance to daptomycin, vancomycin, and dalbavancin. The seesaw effect was most widely observed for cephalexin (6 of 6 strains) and nafcillin (4 of 6 strains), whereas the occurrence of the seesaw effect with meropenem, ceftriaxone, and ceftaroline was more variable and none of the strains evaluated here displayed a clear seesaw effect with cefoxitin.

### Membrane lipids in GP/LP/LGP resistance

Untargeted lipidomic analysis was performed on all study strains using the previously described hydrophilic interaction liquid chromatography-ion mobility-mass spectrometry (HILIC-IM-MS) method for bacterial lipid profiling (44, 66, 67). Principal component analysis (PCA) of the positive and negative mode datasets showed strong agreement between the modes for both N315 and JE2 strains (Supporting Information Figures SI-1 and SI-2, respectively). The N315-derived strains had clear intergroup separation with tight clustering among the biological replicates. Most of the JE2 strains also showed tight clustering of biological replicates and clear intergroup separation with the exception of JE2-Van4, which had greater intra-group variability.

A summary of the major lipids observed in the N315-derived mutant strains is shown in Figure 2 and in tabular format in Supporting Information Table SI-2. In general, levels of LysylPGs were elevated in N315-Dap1 varying from 1.5-5.4 fold although the changes to LysylPGs 31:0 and 33:0 did not show statistical significance (Figure 2A). In N315-Van8, lysylPGs 34:0 and 35:0 were elevated, but to a smaller degree than in N315-Dap1. However, lysylPGs 31:0 was significantly decreased in N315-Van8. The lysylPGs in N315-Dal0.5 had a trend of decrease relative to the levels in the N315 parent strains but the changes were not significant by student’s t-test. LysylPGs 36:0 and 37:0 were not observed in any of the N315-derived strains.

**Figure 2.**
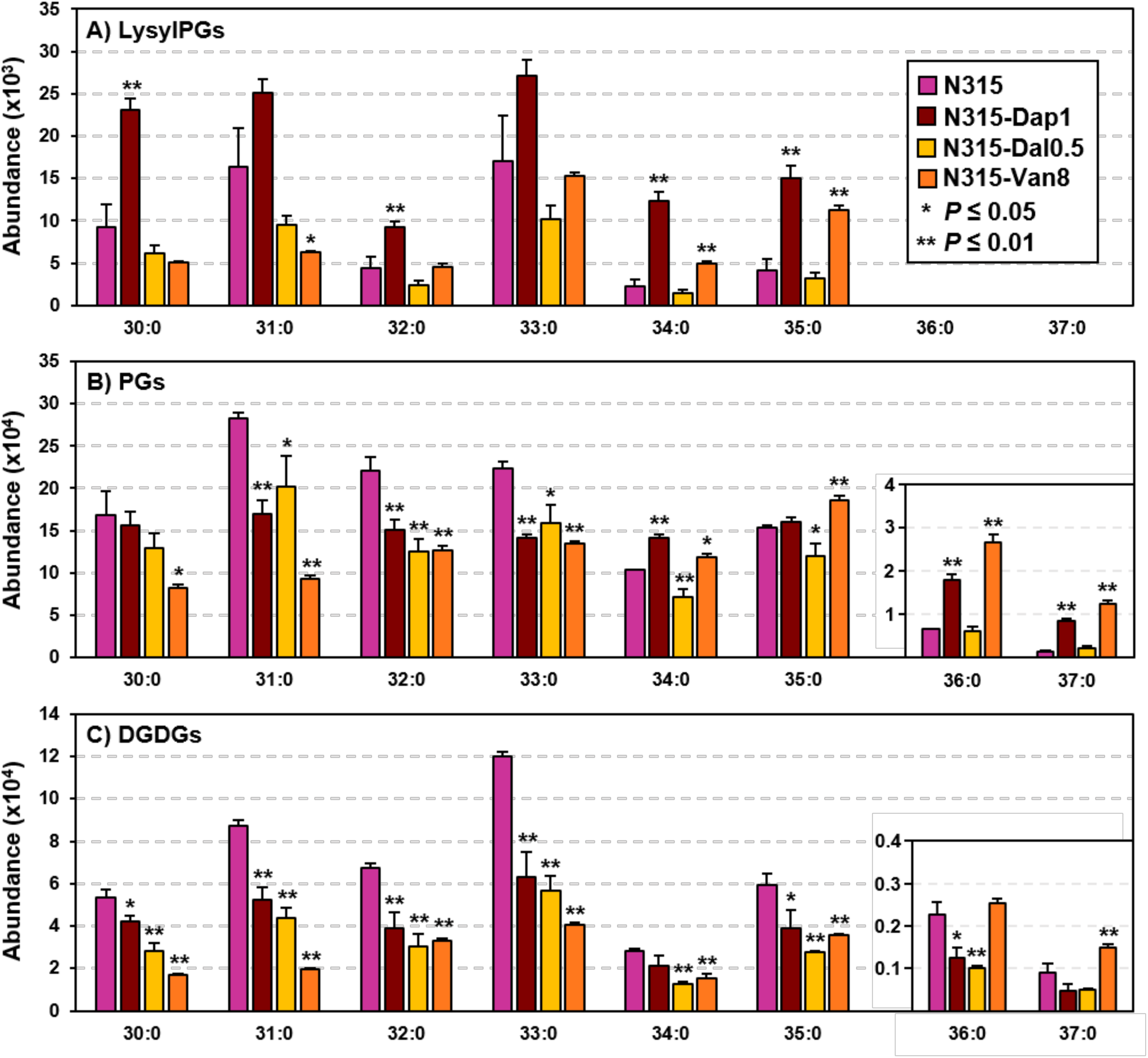
Lipidomics results for *S. aureus* N315 and N315-derived daptomycin-(Dap1), dalbavancin-(Dal0.5) and vancomycin-resistant (Van8) strains. A) Major species of lysyl-phosphatidylglycerols (LysylPGs) from HILIC-IM-MS analysis in positive ionization mode. B) Major species of phosphatidylglycerols (PGs) from HILIC-IM-MS analysis in negative ionization mode. C) Major species of diglucosyl diacylglycerols (DGDGs) from HILIC-IM-MS analysis in positive ionization mode. Error bars represent the standard deviation of the mean. Statistical significance was determined by Student’s t-test *P* ≤ 0.05, two-tailed with equal variance.

Levels of PGs 30:0 to 33:0 (Figure 2B) were decreased in all N315-derived mutant strains relative to parent N315 but change in PG 30:0 is only significant for N315-Van8. On the other hand, long-chain PGs 36:0 and 37:0 were elevated by 4- and 8-fold, respectively, in N315-Van8 relative to parent N315, whereas PGs 34:0 and 35:0 were modestly increased (1.1- and 1.2-fold). A similar trend was observed in N315-Dap1, with elevated levels of 34:0 (1.4-fold), 36:0 (2.7-fold) and 37:0 (5.7-fold).

Major species of DGDGs (Figure 2C) in all N315 derivatives were decreased by 1.3-to 4.5-fold. DGDG 36:0 and DGDG 37:0 were also decreased in N315-Dap1 and N315-Dal0.5 by a similar magnitude. However, DGDG 36:0 and DGDG 37:0 were slightly elevated by 1.1 and 1.6-fold, respectively, in N315-Van8 compared to parent N315.

In JE2, the amount of lysylPGs (Figure 3A, Table SI-3) was higher overall in JE2-Dap2 than in the other JE2-derived strains, similar to those observed in N315-Dap1, with increases from 1.6-fold for lysylPG 30:0 to 5.2-fold for lysylPG 35:0. LysylPGs were also trending higher overall in JE2-Van4 than the parent, but only lysylPGs 34:0 and 35:0 showed statistical significance. In JE2-Dal2, the levels of lysylPGs varied based on the fatty acyl composition: lysylPGs 30:0, 31:0, and 33:0 were decreased whereas lysylPGs 32:0, 34:0 and 35:0 were increased in JE2-Dal2 relative to the parent strain. LysylPGs 36:0 and 37:0 were not detected in any JE2 strains, as in N315 strains.

**Figure 3.**
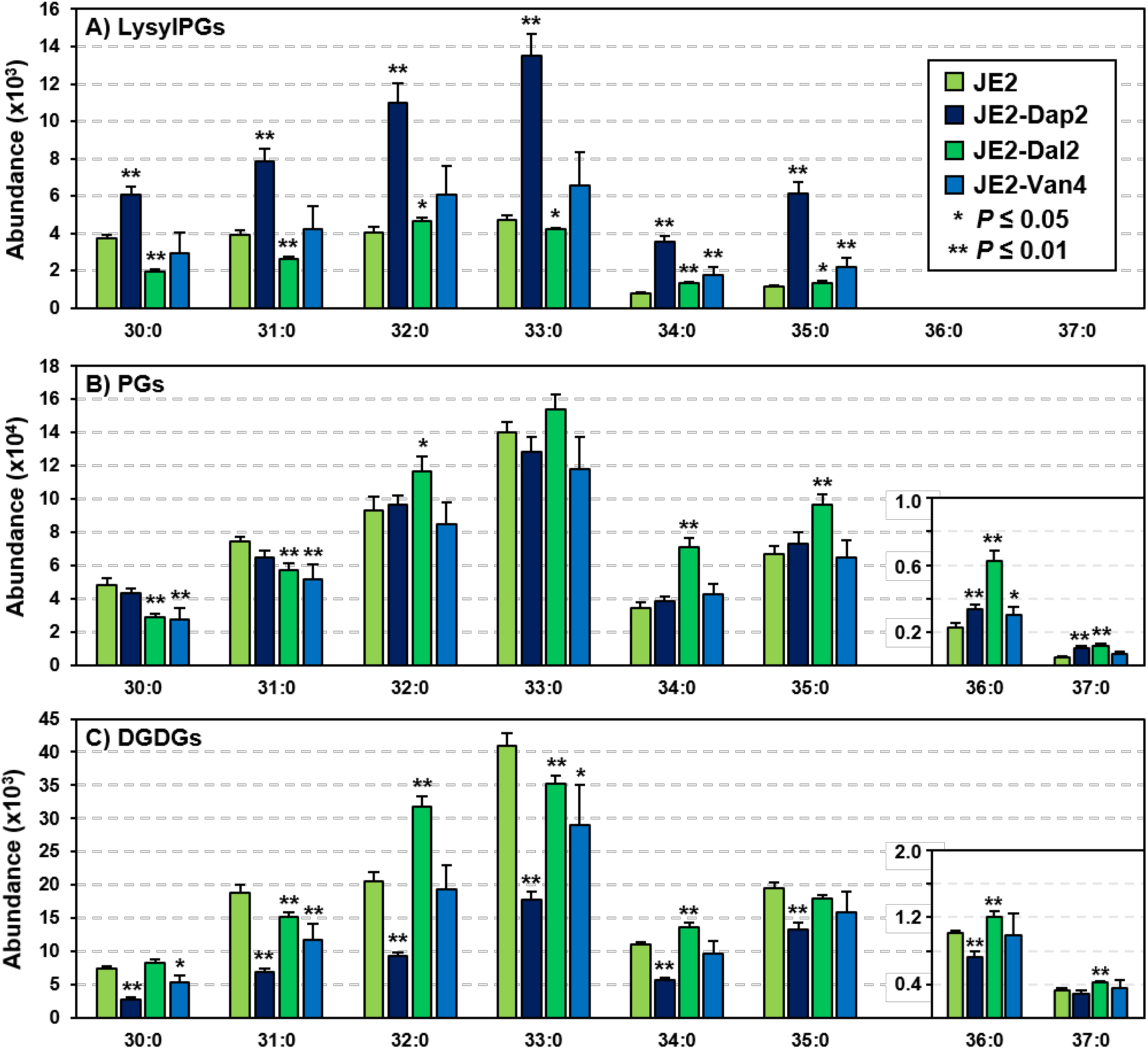
Lipidomics results for *S. aureus* JE2 and JE2-derived daptomycin-(Dap2), dalbavancin-(Dal2) and vancomycin-resistant (Van4) strains. A) Major species of lysyl-phosphatidylglycerols (LysylPGs) from HILIC-IM-MS analysis in positive ionization mode. B) Major species of phosphatidylglycerols (PGs) from HILIC-IM-MS analysis in negative ionization mode. C) Major species of diglucosyl diacylglycerols (DGDGs) from HILIC-IM-MS analysis in positive ionization mode. Error bars represent the standard deviation of the mean. Statistical significance was determined by Student’s t-test *P* ≤ 0.05, two-tailed with equal variance.

For JE2-Dap2, the most abundant species of PGs (Figure 3B, Table SI-3) were unchanged relative to the parent strain, but the long-chain PGs 36:0 and 37:0 were 1.5- and 2-fold higher. In JE2-Dal2, short-chain PGs 30:0 and 31:0 were decreased, whereas PGs 32:0 to 37:0 were elevated to varying degrees. The fold-change increase for the long-chain PG 36:0 and 37:0 (at 2.7- and 2.4-fold, respectively) was greater than that of the other PGs species in JE2-Dal2. In JE2-Van4, only minor decreases in PGs 30:0 and 31:0 and an increase in PG 36:0 relative to the parent strain were observed.

All major species of DGDGs (Figure 3C, Table SI-3) were significantly decreased in JE2-Dap2 by 1.5-to 2-fold. In JE2-Dal2, DGDGs 30:0, 32:0, 34:0, 36:0 and 37:0 were elevated by 1.1-to 1.5-fold relative to the parent JE2 strain, but DGDGs 31:0, 33:0, and 35:0 were decreased by a similar degree. In JE2-Van4, DGDGs 30:0, 31:0, and 33:0 were decreased, whereas DGDGs 32:0, 34:0, 35:0, 36:0 and 37:0 were relatively unchanged.

Free fatty acids (FFA) were also analyzed in all strains. There appeared to be a consistent trend of increased FFAs 14:0, 15:0, 20:0, and 21:0 in the resistant strains relative to their matching parent strain. However, changes to other FFA species varied between the two genetic backgrounds (Figures SI-3&4, Tables SI-2&3).

### Correlation of lipid levels and MICs

Strains of N315 and JE2 in which a strong seesaw effect with nafcillin and cephalexin occurred also displayed higher levels of PG species 34:0, 35:0, 36:0, and 37:0. Pearson correlation coefficients were determined to test the relationship between levels of individual lipid species and antibiotic susceptibility, with separate analyses performed for each lipid class in the N315 set and the JE2 set. The Pearson coefficients, *r*, for the correlation of individual PGs species with MICs for several antibiotics are shown in Figure 4. Additional Pearson correlation results, including Pearson *P* values, can be found in Supporting Information Figures SI-5 through SI-11.

**Figure 4.**
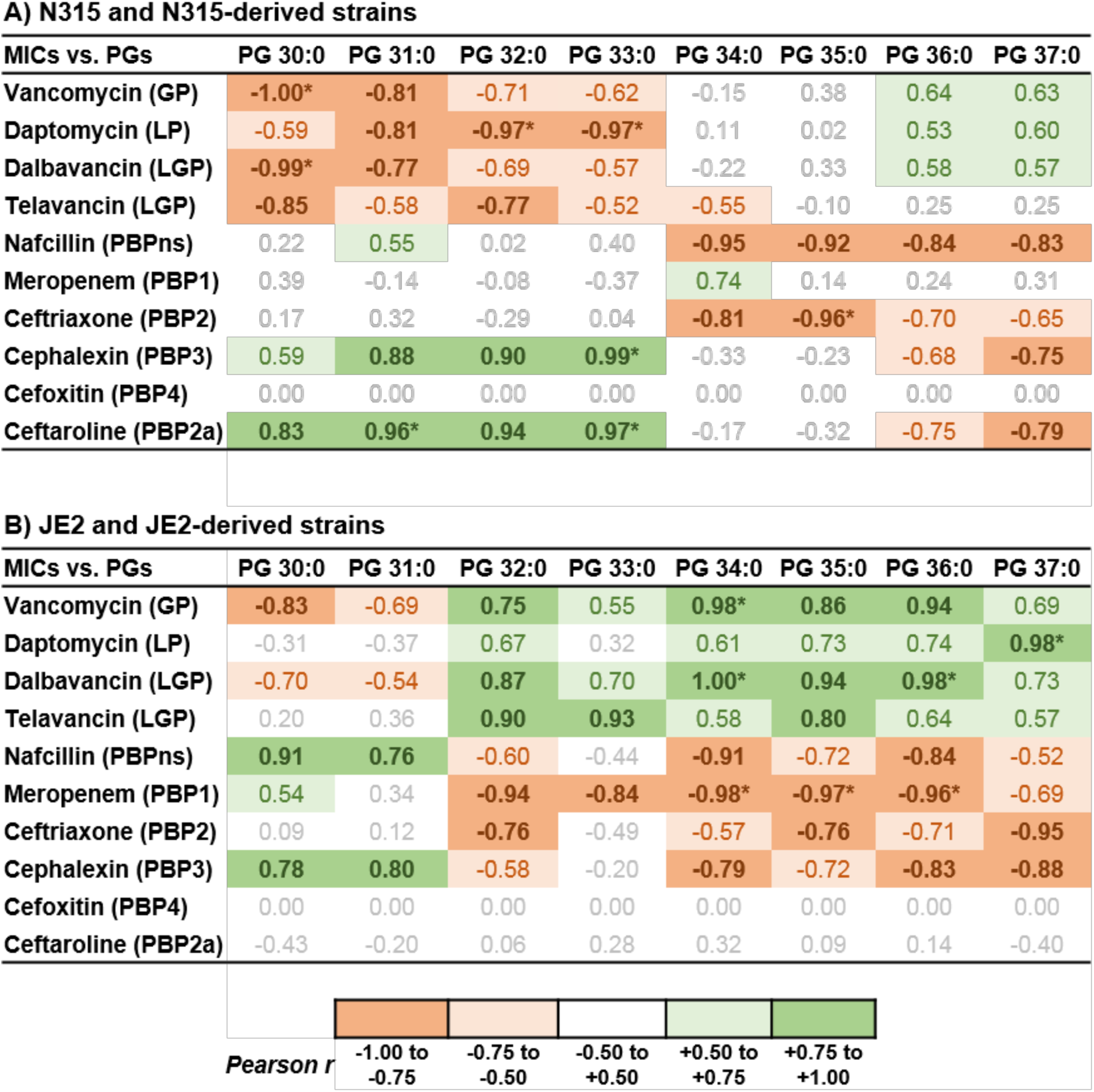
Pearson correlation coefficients, *r*, for the relationship between levels of individual PG species and minimum inhibitory concentrations against various β-lactam and peptide-based antibiotics for A) N315 and N315-derived strains (n=4) and B) JE2 and JE2-derived strains (n=4). *, *P* ≤ 0.05, two-tailed.

In N315 and N315-derived strains, the levels of PGs 30:0 to 33:0 were negatively correlated with the MICs for peptide-based antibiotics vancomycin, daptomycin, dalbavancin, and telavancin. The correlations of PG 30:0 with vancomycin MICs and dalbavancin MICs were particularly strong, with coefficients of −1.0 and - 0.99, respectively, and *P*-values less than 0.05. PGs 32:0 and 33:0 demonstrated a strong negative correlation with daptomycin MICs, with *r* = −0.97 for both and *P*-values less than 0.05. These same PG species demonstrated a positive correlation with MICs of the cephalosporin antibiotics cephalexin and ceftaroline. The positive correlation between PG 31:0 and ceftaroline MICs were also strong (*r* ≥ +0.95, *P* ≤ 0.05). On the other hand, the MICs of nafcillin and ceftriaxone were negatively correlated with the levels of PGs 34:0, 35:0, 36:0 and 37:0. PGs 36:0 and 37:0 were also negatively correlated with the MICs of additional beta-lactams, cephalexin and ceftaroline.

In JE2, the vancomycin and dalbavancin MICs were negatively correlated with the levels of PGs 30:0 and 31:0 but did not show as strong of a correlation as was observed for the N315-derived strains. PGs 30:0 and 31:0 were positively correlated with nafcillin and cephalexin MICs. In general, the MICs for the peptide-based antibiotics were positively correlated with the levels of PGs 32:0 to 37:0. The correlations of vancomycin MICs with PG 34:0, daptomycin MICs with PG 37:0, and dalbavancin MICs with PGs 34:0 and 36:0 were very strong (*r* ≥ +0.95, *P* ≤ 0.05). The MICs of nafcillin, meropenem, ceftriaxone, and cephalexin were in general negatively correlated with the levels of PGs 32:0, 34:0, 35:0, 36:0 and 37:0. The Pearson coefficients for the relationships between meropenem and PGs 34:0 to 36:0 demonstrated a strong and significant negative correlation (*r* ≤ −0.95, *P* ≤ 0.05).

Correlation analysis between MICs and DGDGs, lysyPGs and FFAs was also performed. Although there were significant correlations between DGDGs and various MICs in N315-derived strains, these correlations were more sporadic and less significant in the JE2-derived strains (Figures SI-6&7). Coefficients from the correlation analysis of lysylPGs vs. MICs and FFAs vs. MICs were in general less significant and varied more between the two genetic backgrounds (Figures SI-8&9). These observations suggest that DGDGs, lysylPGs, and FFAs are not reliable indicators of susceptibility against GP/LP/LGP or beta-lactams across different genetic backgrounds.

## DISCUSSION

In this study, varying degrees of beta-lactam seesaw effect was observed among the vancomycin-, daptomycin- and dalbavancin-resistant mutants derived from MRSA strains N315 and JE2 by serial passage, depending both on the resistance profile of the strain and the beta-lactam antibiotic used. The beta-lactam antibiotics used to evaluate the seesaw effect were chosen based on their specificities for binding different PBPs. The beta-lactams showing the largest and most consistent seesaw effect with daptomycin, vancomycin, and dalbavancin were the PBP-nonspecific anti-staphylococcal penicillin, nafcillin, and the PBP3-specific cephalosporin, cephalexin. However, a more limited seesaw effect, both in terms of susceptibility improvement and number of observations, was also observed with ceftriaxone (PBP2), ceftaroline (PBP2a) and meropenem (PBP1).

The occurrence of the seesaw effect has been most widely explored for oxacillin (PBP-nonspecific) and ceftaroline in daptomycin-resistant (Dap-R) MRSA and VISA. The seesaw effect between daptomycin and oxacillin was demonstrated *in vitro* by Yang et *al.* (16) and *in vivo* by Lee *et al.* (17), who described a case study where high-dose daptomycin therapy induced daptomycin-resistance but improved oxacillin susceptibility. Mehta *et al.* (58) noted the occurrence of the seesaw effect with both oxacillin and nafcillin in daptomycin-resistant MRSA with fold-change increases in susceptibility comparable to our observations with nafcillin in N315-Van8 and N315-Dap1. In a study of 150 MRSA strains, Barber *et al.* (11) showed that ceftaroline MICs were negatively correlated with daptomycin and vancomycin MICs as would be expected in the occurrence of the seesaw effect. In a similar study of 153 hVISA and VISA isolates, vancomycin MICs were significantly inversely correlated with oxacillin, cefoxitin, and ceftaroline MICs (10). We have also (19) observed that even when there is no change in beta-lactam MIC as vancomycin or daptomycin MICs increase, increased susceptibility to ceftaroline can be measured by population analysis profiling and in pharmacodynamic response in *in vitro* PK/PD models. Our present findings with ceftaroline display modest seesaw effect with vancomycin and dalbavancin in the N315 genetic background and with daptomycin in both the N315 and JE2 backgrounds.

The differential expression of PBPs in DAP-R and VISA strains is thought to be the source of the beta-lactam seesaw effect. VISA strains have been shown to have reduced PBP4 and PBP2a and are particularly susceptible to beta-lactams targeting PBP2, which is presumed to take over cell wall synthesis in the absence of PBP2a and PBP4 (14, 68). Berti *et al.* (59) observed that the MICs for PBP1-specific antibiotics, such as meropenem, were decreased 2-to 4-fold in a clinical isolate of MRSA with daptomycin resistance. This led to the exploration of the role of PBP1 in daptomycin resistance, where it was found that expression of the *pbpA* transcript, which encodes PBP1, was increased upon daptomycin exposure while transcription of genes encoding for PBP2, PBP3 and PBP4 remained unchanged. This study also observed increased resistance to daptomycin when pbpA transcription was increased in a strain of MRSA, COL, with *pbpA* under an inducible promotor. However, we did not observe the seesaw effect with meropenem in either of the strains with reduced daptomycin susceptibility. Rather, we have observed more widespread seesaw effect for PBP3-targteting cephalexin and PBP-nonspecific nafcillin.

While named for their interaction with beta-lactam antibiotics, the primary function of penicillin binding proteins is the synthesis of cell wall peptidoglycan and alteration in the activity or expression of PBPs has measurable effects on the cell wall. A common, but not universal, phenotype for both daptomycin-resistant MRSA and VISA is a thicker cell wall, which serves to prevent daptomycin and vancomycin from reaching their targets of the cell membrane or the cell wall-cell membrane interface, respectively (41, 50, 69). The first study to demonstrate a seesaw effect between vancomycin and beta-lactams noted that there was substantially less cross-linking in the cell wall of the highly vancomycin-resistant strain (12). It has been proposed that this decreased cross-linking contributes to the thickened cell walls of VISA as a result of decreased PBP4 levels or activity (14), although this has not been observed universally (19). Several studies have shown that exposure of Dap-R or VISA to beta-lactam antibiotics results in a thinning of the cell wall. Ceftaroline treatment of VISA with daptomycin resistance reduced the thickness of the cell wall by approximately 9 nm (57). In Dap-R MRSA with seesaw effect with PBP1-specific beta-lactams, the cell wall thickness was increased, rather than decreased, after overnight exposure to PBP1-specific beta-lactams (59). However, exposure to the PBP3-specific cephalosporin cefaclor did result in a thinning of the cell wall.

The two-component system for cell wall stress, VraSR, is often associated with resistance to antibiotics that target the cell wall, including methicillin, vancomycin, and daptomycin (70–72), and was identified as a frequently mutated target in our antibiotic selected strains. Renzoni *et al.* have proposed that the reduced levels of PBP2a in daptomycin-resistant MRSA with beta-lactam susceptibility is driven by the *vraSR* system (20). Several studies have observed that overexpression of the *vraSR* system in Dap-S MRSA simultaneously decreased daptomycin susceptibility and increased susceptibility to oxacillin (20, 58). Boyle-Vavra *et al.* (73) and McCallum *et al.* (74) have independently verified that *vraSR* is under the regulation of the upstream gene *vraT*, and mutations in *vraT* have accordingly been identified in VISA and dalbavancin nonsusceptible VISA isolates (38, 54). Here, we identified mutations in *vraT* in the both strains with reduced susceptibility to dalbavancin (N315-Dal0.5, JE2-Dal2), as well as in JE2-Van4, and a mutation in *vraS* in N315-Dap1 (Table 1). However, the presence of mutations within the *vraTSR* system did not confer improved beta-lactam susceptibility universally. These results suggest that occurrence of the seesaw effect is mediated by other mechanisms.

The phenomenon of decreased negative cell surface charge in Dap-R is often mediated through reduced membrane levels of the negatively-charged PGs, which can be achieved through several mechanisms. The most widely associated mutation associated with the daptomycin resistance phenotype is that of *mprF* (45, 46, 48, 49). Gain-of-function mutation of this gene results in increased lysinylation of PGs and effectively transforms a negatively-charged lipid into one with a net-positive charge. Many studies of daptomycin-resistant pathogens have confirmed this phenotype through lipidomics analyses (40, 41, 75, 76). The JE2-Dap2 strain characterized here contained a mutated *mprF* gene and showed substantially increased levels of lysylPGs, but these changes did not occur at the expense of reduced PG levels. The N315-Dap1 strain also had modestly increased levels of lysylPGs and decreased levels of the major PG species in the absence of changes to the *mprF* coding sequence. Rather, the alteration of membrane lipids in this strain could be attributed to the mutations in *vraS* and in *yycG*, the histidine kinase of the *yycFG* two-component system involved in cell membrane stress response and fatty acid biosynthesis (77, 78). In agreement with our previous findings in a highly daptomycin-resistant strain of MRSA (44), the amount of minor long-chain PG species, PG 36:0 and 37:0, were significantly elevated in both N315-Dap1 and JE2-Dap2.

Although PGs 36:0 and 37:0 are typically minor PGs in MRSA, it is plausible that the increase in PGs with long chain fatty acyl tails may have an impact on the fluidity of the cell membrane and the distribution of membrane-anchored proteins around the cell. As the point for cell division, the majority of cell wall synthesis machinery and, therefore, the majority of newly-synthesized peptidoglycan is localized to the septum in *S. aureus*. Renzoni *et al.* have shown that peptidoglycan insertion is delocalized away from the division septum in daptomycin-susceptible MRSA that has been exposed to daptomycin (20). However, DAP-R MRSA maintains peptidoglycan insertion at the septum after exposure to daptomycin. This suggests that the disruption of the regular membrane organization upon daptomycin exposure also affects the localization of the machinery that is required for cell wall synthesis, such as PBPs. Alternatively, daptomycin-resistant MRSA can resist this daptomycin-induced membrane reorganization and therefore the cell wall synthesis machinery remains at the division septum. We propose that the shift in PG composition towards longer chain fatty acyl tails prevents disruption of the membrane stability by daptomycin so that cell wall synthesis machinery remain localized at the division septum, and that a similar mechanism is occurring in vancomycin- and dalbavancin-resistant MRSA as well.

The exact mechanism behind the seesaw effect is yet to be elucidated, and the mechanism by which a change in membrane composition contributes to beta-lactam susceptibility is unclear at this time. Renzoni *et al.* have proposed that the mprF-mediated increase in lysylPGs in daptomycin resistance prevents the necessary lipidation of the protein, PrsA, involved in the post-translocational extracellular folding of secreted proteins, including PBP2a (20, 79). Without a properly functioning PrsA, the amount of fully-matured PBP2a would be reduced. Lipidation of PrsA is performed by Lgt using the diacylglycerol portion of PG (80). It is possible that the increase in long chain fatty acyl PGs contributes to the dysfunction of PrsA, as proposed by Renzoni *et al.*, if they can no longer function as substrates for Lgt.

To our knowledge, this work represents the first evaluation of membrane lipid composition in VISA and dalbavancin-nonsusceptible MRSA, as well as the first study to explore a relationship between membrane lipids and seesaw effect with beta-lactams. We have demonstrated here that modulation of membrane lipid content is widely observed among MRSA with resistance to vancomycin and dalbavancin in a manner that is not solely attributed to the mutation frequently associated with daptomycin resistance. Among the six mutant strains evaluated in this study, the seesaw effect was observed to at least one, but often several, of the beta-lactams that were tested. Our results provide compelling evidence that: i) membrane lipid composition is implicated in resistance to vancomycin and dalbavancin in addition to daptomycin; ii) the alteration of membrane lipid composition in resistance occurs in a manner that is dependent upon their fatty acyl composition; iii) the abundance of certain short-chain fatty acyl PGs is correlated with cross-resistance to GP/LP/LGP; and iv) the abundance of certain long-chain fatty acyl PGs is correlated with the occurrence of the beta-lactam seesaw effect. Specifically, we found consistent negative correlations between PGs 36:0 and 37:0 with the MICs of nafcillin, ceftriaxone and cephalexin in both genetic backgrounds. As the factors contributing to the seesaw effect continue to be explored, the contribution and significance of membrane lipids to beta-lactam, GP, LP and LGP resistance should not be overlooked.

## MATERIALS AND METHODS

### Reagents

LC/MS grade solvents (water, acetonitrile, chloroform, and methanol) and ammonium acetate were purchased from Thermo Fisher Scientific. Standards for lipidomics were purchased from Avanti Polar Lipids and Nu-Chek prep and prepared as described previously (44, 66). All media were purchased from Thermo Fisher Scientific. Daptomycin and dalbavancin were purchased commercially from their manufacturers (Merck, Allergan) and all other antimicrobials were purchased from Sigma.

### In vitro selection of resistance

The serial passage technique was used for the *in vitro* selection of daptomycin (Dap), dalbavancin (Dal) and vancomycin (Van) non-susceptible mutants from the laboratory *Staphylococcus aureus* strains N315 and JE2 (BEI Resources) in a manner similar to previously described methods (44, 81). Cultures of N315 and JE2 were prepared in Brain Heart Infusion (BHI; Thermo Scientific) broth that was supplemented with 50 mg/L of elemental calcium (BHI-50) for selection with daptomycin and 0.002% polysorbate 80 (BHI-PS80) for selections with dalbavancin to reduce nonspecific binding of dalbavancin to the plastic. Selections against vancomycin were performed in BHI without additional supplementation. Cultures of JE2 and N315 were exposed to 1X the minimum inhibitory concentration (MIC) of the antibiotic, as shown in Table 1, and incubated at 37 °C with shaking at 180 rpm. Visible growth was diluted 1:100 in fresh broth that was supplemented with Dap, Dal or Van at either 2X the previous concentration or 2 µg/mL higher than the previous concentration. This process was repeated until the MICs in Table 1 were achieved.

### Antibiotic susceptibility testing

Each resulting isolate from the serial passage mutant selection was stocked in glycerol tubes, stored at −80ºC, and subcultured on TSA prior to susceptibility testing by broth microdilution in Mueller-Hinton broth (MHB; Becton Dickinson). For the final isolates, susceptibility testing was performed against a collection of beta-lactams antibiotics with different specificities for penicillin binding proteins (PBPs), including nafcillin (PBP non-specific) and cephalexin (PBP3) as shown in Table 1 and meropenem (PBP1), ceftriaxone (PBP2), cefoxitin (PBP4), and ceftaroline (PBP2a) as shown in SI Table 1. Susceptibility against the lipoglycopeptide antibiotic telavancin was also determined (SI Table 1).

### Whole genome sequencing

DNA from parental strains and antibiotic-selected derivatives was extracted using the Ultraclean microbial DNA isolation kit (Mo Bio). Sequencing libraries were prepared as described elsewhere (82, 83), with sequencing performed using an Illumina MiSeq (Illumina, San Diego, CA, USA) with 150-bp paired-end chemistries. Sequence analysis was performed to identify single nucleotide mutations and insertion and deletion mutations in coding sequences as previously (PMID 29089424), against reference genomes of N315 (GenBank Accession BA000018) or JE2 (GenBank Accession CP020619) as appropriate. Variants present in the parental strains were disregarded. Sequence variants were annotated using SnpEFF (84).

### Sample collection and lipid extraction

For the lipidomics analysis, each strain was streaked out onto Brain Heart Infusion agar (BHIA) and allowed to grow overnight at 37 °C. From these plates, inoculums were prepared in sterile water to 2.0 McFarlands and 250 µL was added to 50 mL tubes containing 25 mL of BHI broth supplemented with 50 mg/L of elemental calcium (BHI-50). The same inoculum was used to prepare three replicate cultures for each strain. The cultures were grown overnight at 37 °C with shaking at 180 rpm. Overnight cultures were centrifuged for 10 min at 2,500 x *g* to pellet the bacteria and 20-22 mL of supernatant was aspirated from the tube. The pellet and 3-5 mL of supernatant were transferred to pre-weighed 10 mL glass centrifuge tubes and centrifuged. The remaining supernatant was aspirated, and the pellets were dried in a vacuum concentrator. The average dried pellet weights per strain were: N315, 10.9 ± 1.4 mg; JE2, 34.6 ± 4.0 mg. Samples were stored at −80 °C until analysis.

Bacterial lipids were extracted using the method of Bligh and Dyer, as described previously (44). Briefly, 1 mL of water was added to the pelleted and dried bacteria. The resulting suspensions were sonicated in an ice bath for 30 min to dislodge the dried pellets and homogenize the suspension. A chilled solution of methanol and chloroform (2:1 v/v, 4 mL) was added to each tube, followed by 5 min of mixing and the addition of 1 mL of chilled chloroform and 1 mL of chilled water. The samples were vortexed briefly and centrifuged for 10 min at 10 °C and 1,000 x *g* to separate the organic and aqueous layers. The organic layers were collected into clean 10 mL glass centrifuge tubes and dried in a vacuum concentrator. The dried lipid extracts were reconstituted in 500 µL of 1:1 chloroform/methanol. Based on the dried pellet weights, 5-10 µL of the lipid extract was transferred to an LC vial, dried under Ar, and reconstituted to 200 µL with 2:1 acetonitrile/methanol.

### Liquid chromatography

Chromatographic separation of bacterial lipids was performed as described previously (44, 66). Briefly, hydrophilic interaction liquid chromatography (HILIC) was performed with a Phenomenex Kinetex HILIC column (2.1×100 mm, 1.7 µm) maintained at 40 °C on a Waters Acquity FTN UPLC (Waters Corp., Milford, MA, USA) operating at a flow rate of 0.5 mL/min. The solvent system consisted of: A) 50% acetonitrile/50% water with 5 mM ammonium acetate; and B) 95% acetonitrile/5% water with 5 mM ammonium acetate. The linear gradient was as follows: 0-1 min, 100% B; 4 min, 90% B; 7-8 min, 70% B; 9-12 min, 100% B. A sample injection volume of 5 µL was used for all analyses.

### Ion mobility-mass spectrometry

The Waters Synapt G2-Si HDMS platform was used for lipidomics analysis (44, 66). Effluent from the UPLC was introduced through the electrospray ionization (ESI) source. ESI capillary voltages of +2.5 and −2.0 kV were used for positive and negative analyses, respectively. Additional ESI conditions were as follows: sampling cone, 40 V; extraction cone, 80 V; source temperature, 150 °C; desolvation temperature, 500 °C; cone gas, 10 L/hr; desolvation gas, 1000 L/hr. Mass calibration over *m/z* 50-1200 was performed with sodium formate. Calibration of ion mobility measurements was performed as previously described (85). IM separation was performed with a traveling wave height of 40 V and velocity of 500 m/s. Data was acquired for *m/z* 50-1200 with a 1 sec scan time. Untargeted MS/MS was performed in the transfer region with a collision energy ramp of 35-45 eV. Mass and drift time correction was performed post-acquisition using the leucine enkephalin lockspray signal.

### Data analysis

Data alignment, peak detection and normalization was performed in Progenesis QI (Nonlinear Dynamics) based on the chromatographic region from 0.4 to 9.0 min. A pooled quality control sample was designated as the alignment reference. The default “All Compounds” method of normalization was used to correct for variation in the total ion current amongst samples. For initial statistical analyses, the resulting features were filtered by ANOVA *P* ≤ 0.05 for the JE2 analysis, and *P* ≤ 0.001 and fold-change ≥ 2 for the N315 analysis. Student’s t-tests for two samples were performed using a two-tailed distribution and equal variance. Lipid identifications were made based on an in-house version of LipidPioneer modified to contain the major lipid species observed in *S. aureus*, including free fatty acids (FFAs), diglucosyl diacylglycerols (DGDGs), phosphatidylglycerols (PGs), and lysyl phosphatidylglycerols (LysylPGs) with fatty acyl compositions ranging from 25:0 to 38:0 (total carbons: total degree unsaturation) (44, 86). Identifications were made within 15 ppm mass accuracy.

### Data availability

Whole genome sequencing data from this study are available from the NCBI Sequence Read Archive (SRA; http://www.ncbi.nlm.nih.gov/sra) under BioProject number <u>PRJNA547605</u>.

## ACKNOWLEDGEMENTS

This study was supported by a grant from the University of Washington School of Pharmacy Faculty Innovation Fund and University of Washington Royalty Research Fund (A128444) to LX and BJW, the startup fund to LX from the Department of Medicinal Chemistry in the School of Pharmacy at the University of Washington, and the National Institute of Allergy and Infection Diseases R21AI132994 and R01AI136979 to BJW and LX. BJW has received research grants from commercial sources, including Shionogi.

## AUTHOR CONTRIBUTIONS

BJW and LX conceived the study. KMH and LX designed and performed the lipidomics experiments and analyzed the mass spectrometry data. BJW, KMH, TS and NA performed the resistance passage studies and other microbiological aspects of the experiments. SJS, AW, EAH, KP, and KM performed whole genome sequencing and analysis. All authors reviewed the data, prepared the manuscript, and approved its final version.

